# Comparative transcriptome analysis in contrasting finger millet (*Eleusine coracana* (L.) Gaertn) genotypes for heat stress

**DOI:** 10.1101/2023.03.19.533344

**Authors:** Etika Goyal, Singh Kumar Amit, Mahesh Mohanrao Mahajan, Kumar Kanika

## Abstract

*Eleusine coracana* (L.) Gaertn is a crucial C_4_ species renowned for its stress robustness and nutritional significance. Because of its adaptability traits, finger millet (ragi) is a storehouse of critical genomic resources for crop improvement. However, more knowledge about this crop’s molecular responses to heat stress must be gained. Hence, in the present study, we generated RNA seq data from the leaf tissue of the finger millet to observe the physiological changes and gene expression study in heat-sensitive (KJNS-46) and heat-tolerant (PES-110) genotypes of Ragi in response to high temperatures. On average, each sample generated about 24 million reads. Nearly 684 transcripts were differentially expressed (DEGs) between the heat-stressed samples of both genotypes. Pathway analysis and functional annotation analysis revealed the activation of various genes involved in response to stress, precisely heat, oxidation-reduction process, water deprivation, heat shock protein and transcription factors, calcium, and kinase signaling. The basal regulatory genes, such as bZIP, were involved in response to heat stress, indicating that heat stress activates genes related to basal regulatory processes or housekeeping. A substantial percentage of the DEGs belonged to proteins of unknown functions (PUFs), i.e., uncharacterized. The expression pattern of a few selected DEGs genes was analyzed in both genotypes by quantitative RT-PCR. The present study found some candidate genes and pathways that may confer tolerance to heat stress in ragi. These results will provide valuable information to improve heat tolerance in heat-susceptible agronomically important varieties of ragi and other crop plants.

## 1. Introduction

Finger millet is an allotetraploid (2n = 4x = 36), a member of the family Poaceae and sub-family Chloridiodeae (Goron and Raizada 2015). It was introduced into the Western Ghats of India around 3000 BC. Consequently, India became the secondary center of diversity for finger millet (Hittalmani et al. 2017). It is rich in amino acids like calcium, methionine, and iron, which are lacking in the diet of the poor, who live on starchy staples such as cassava, rice, maize, etc. Finger millet species have some enduring characteristics, such as tolerance to heat, cold, drought, etc. (Keller and Seehausen 2012).

Heat stress activates HSFs (heat shock transcription factors), proteins (HSPs), and various metabolic pathways, which help plants obtain basal and acquired thermo-tolerance (Baniwal et al. 2014). The heat stress response is complex regulatory networks that involve large-scale gene reprogramming at the transcriptome level (Liu et al. 2013; Dittami et al. 2009). Transcriptomic studies under heat stress have been reported in grape, pepper, spinach, switchgrass, tomato, cotton, cucumber, *Arabidopsis*, and jujube (Liu et al. 2012; Li et al. 2015; Yan et al. 2016; Li et al. 2016; Frank et al. 2009; Wang et al. 2012; Qi et al. 2012; Yasuhiro et al. 2015; Juan et al. 2021). Identifying genes associated with response to heat stress can significantly facilitate the development of improved cultivars, which can be done with large-scale genomic resource availability. One of the significant drawbacks in this direction is the need for finger millet genomic resources. Transcriptome analysis of finger millet under salinity, drought stress, and developing spikes has been carried out in the past (Kumar et al. 2015; Gururani et al. 2020; Rahman et al. 2014; Hittalmani et al. 2017; Li et al. 2021; Parvathi et al. 2019).

Unfortunately, finger millet has never been studied for heat stress response; therefore, comparative differential gene expression analysis can improve our understanding of the molecular mechanisms in contrasting genotypes for tolerance towards heat stress. This study will also shed light on adaptation to local climates, especially at the transcriptome level. It will elucidate the complex mechanisms and improve our understanding of the response to heat stress. This, in turn, will help us fine-tune our efforts for future crop improvement programs. This will also enable us to predict the response of this nutritionally rich crop to changes in environmental conditions due to climate change in the future.

## 2. Material and Methods

### 2.1 Plant Material and Heat Treatment

Contrasting finger millet genotypes, i.e., PES-110 (heat tolerant) and KJNS-46 (heat susceptible), were procured from GKVK, UAS, Bangalore, Karnataka. All plants were germinated and maintained at 28°C for three weeks (12h light/10h dark) at the National Phytotron Facility, IARI Campus, New Delhi. The heat treatments were initiated in the middle of the day, namely 7hr after light onset: a) short-term (T1): 42°C treatment up to 4hrs; and b) long-term: 42°C treatment for 4hrs for three continuous days. To reduce side effects, the cabinet was near the control plants. Leaf samples from both sets were immediately collected and flash-frozen in liquid N_2_ and stored at - 80°C until further use.

### 2.2 Physiological measurements

Physiological parameters, including membrane stability index (MSI) and proline were measured following standard methods in three different samples in each treatment for each genotype to analyze the physiological changes under heat stress before the transcriptome study (Mahajan et al. 2017).

### 2.3 RNA sequencing

RNA extraction and purification were done using RaFlex™ Total RNA Isolation Kit (GeNei^™^, Bangalore, India). Equimolar amounts of T1 and T2 RNA samples were pooled. Commercial service providers (NxGenBio Life Sciences, New Delhi, India) did the library preparation and sequencing on Illumina HiSeq 2000 instrument using 2×100bp paired-end chemistry. The phenotype towards heat stress of the two accessions (tolerant/sensitive) along with conditions (control/heats stress) were used as sample names (for example, KJNS-C and KJNS-T refer to KJNS-46 heat susceptible-control and KJNS-46 heat susceptible-treatment, respectively). FastQC was used to control the data quality, and Trinity *de novo* assembler assembled the reads into contigs (Grabherr et al. 2011). Bowtie aligner aligned clean reads back to the assembled non-redundant reference transcriptome (Langmead et al. 2009). The resulting quantification was expressed as FPKM (Fragments per Kilobase of Transcript per Million Mapped Reads). For calculating the differential expression of genes among the control vs. treatment samples (PES-C vs. PES-T; KJNS-C vs. KJNS-T) and PES-T vs. KJNS-T, the RSEM package was used with default parameters.

### 2.4 Data analysis

Blast homology searches and sequence annotations were done using the Blast2GO tool v.4.0. The assembled sequences were compared against the NCBI non-redundant (nr) protein database via Blast X, using an E-value cut-off of 1.0E^-3^. Gene ontology (GO) terms were assigned to the unigenes, and the unigenes were also aligned to the COG database to predict and classify proteins. KEGG pathways were assigned to the assembled sequences using the online KEGG Automatic Annotation Server (KAAS). For the identification of transcription factors in finger millet transcriptome, all the assembled unigenes were analyzed against the PlantTFcat online tool (Goyal et al. 2016a).

### 2.5 Validation by qRT-PCR

Twelve expressed genes were validated using the qRT-PCR method, and *actin* was used as a reference for gene expression normalization. RT-PCR was carried out as mentioned by Goyal et al. 2016a. The gene-specific primers are listed in Supplementary Table S1.

## 3. Results

### 3.1 Physiochemical variations in response to heat stress of two genotypes

To confirm the difference in heat tolerance between the heat-tolerant and sensitive genotypes of ragi, we measured the MSI and proline content as mentioned in the Material and Method section. Compared with the control treatment, there was no significant difference in the MSI content in ‘PES-110’ leaves after 3hr or one day of heat stress in both treatments. But exposure to heat stress for three days and longer hours significantly increased the MSI content (data not shown). Exposure to heat stress significantly increased the proline content in the leaves of ‘KJNS-46’ and ‘PES-110’ in both treatment sets compared with the control treatment (data not shown).

### 3.2 Transcriptome sequencing and *de-novo* assembly

The control and heat stress leaf tissues of KJNS-46 and PES-110 ragi genotypes were used for RNA isolation and sequencing. Approximately 24 million reads were generated from each sample, leading to 145.5 million pair-end reads obtained from the four tissue samples. The raw transcriptome data have been deposited at NCBI as Sequence Read Archive (SRA), with the accession numbers SRR3156038, SRR3156044, SRR3146431, and SRR3146438 for PES-C, PES-T, KJNS-C, and KJNC-T, respectively.

The assembly generated 58,430, 87,046, 65,866, and 80,131 transcripts in KJNS-C, KJNS-T, PES-C, and PES-T, respectively. Approximately 51% of these contigs had a length ≥500 bp. The summary of the sequence assembly statistics is given in Table 1. The average GC content of finger millet was 48.92%, which is in the range of GC levels of monocot coding sequences (Goyal et al. 2016 a, b; Mahajan et al. 2017, 2020). A total of 13,642, 27,910, 17,031, and 23,659 ORFs were detected in KJNS-C, KJNS-T, PES-C, and PES-T, respectively (Table 1). In KJNS-C and KJNS-T, the longest ORF was 3,234 and 1,711nt, respectively, whereas, in PES-C and PES-T, this was 1,948 and 2,336nt, respectively. The length distribution of contigs is shown in Supplementary Fig. 1A. Proteome analysis of total transcripts of all four samples against the rice and foxtail millet databases revealed that, in total, 39,507 and 40,758 transcripts of KJNS-C; 54,103 and 55,197 transcripts of KJNS-T; 44,161 and 45,520 transcripts of PES-C and 49,960 and 51,258 transcripts of PES-T showed similarity with them, respectively (Supplementary Fig. 1B).

**Table 1:** Annotation statistics of finger millet transcriptome under heat stress. ORF, open reading frame

| Properties | KJNS-C | KJNS-T | PES-C | PES-T |
| --- | --- | --- | --- | --- |
| Total reads | 22,705,966 | 23,259,542 | 24,249,804 | 25,998,907 |
| Total Bases | 4,563,899,166 | 4,675,167,942 | 4,874,210,604 | 5,225,780,307 |
| Total No. of Contigs (n) | 58,430 | 87,046 | 65,866 | 80,131 |
| Minimum contigs length (bp) | 201 | 201 | 201 | 201 |
| Maximum contigs length (bp) | 7,242 | 10,546 | 7,222 | 10,148 |
| Average contigs length (bp) | 744.71 | 899.09 | 786.19 | 844.65 |
| GC content (%) | 49.20 | 48.49 | 49.32 | 48.95 |
| ORF found | 13,642 | 27,910 | 17,031 | 23,659 |
| N50 | 1,178 | 1,540 | 1,258 | 1,407 |

In the plant ref database, out of 60,549 assembled unigenes in KJNS-46, only 54,565 had at least one significant match, and 32,890 got a UniProt ID. However, for PES-110, out of 55,864 assembled unigenes, 50,854 unigenes showed homology against the plant ref database, whereas; only 31,118 unigenes had UniProt ID (Supplementary Table S2 A, B).

Mapping all the reads onto the nr dataset revealed that the number of reads corresponding to each transcript ranged from 0 to 11,540 for KJNS-C, 67,838 for KJNS-T, 14,949 for PES-C, and 123 FPKM for PES-T. This indicates a wide range of expression levels of finger millets transcripts in response to heat stress. It also indicated that even the low-expressed transcripts are detected in the present analysis.

### 3.3 Transcripts expressed under treatment conditions in both the genotypes

In the PES-110, out of 55,864 unigenes, 1,972 unigenes were differentially expressed. Out of this, 1,044 were up-, whereas only 928 unigenes were down-regulated in PES-T compared to PES-C. In the KJNS-46, out of 60,549 unigenes, 2,637 unigenes were differentially expressed. Out of this, 1,068 were up-, whereas 1,569 unigenes were down-regulated in KJNS-T compared to KJNS-C. A Venn diagram representing the differentially expressed genes is shown in Fig. 1.

**Fig 1.**
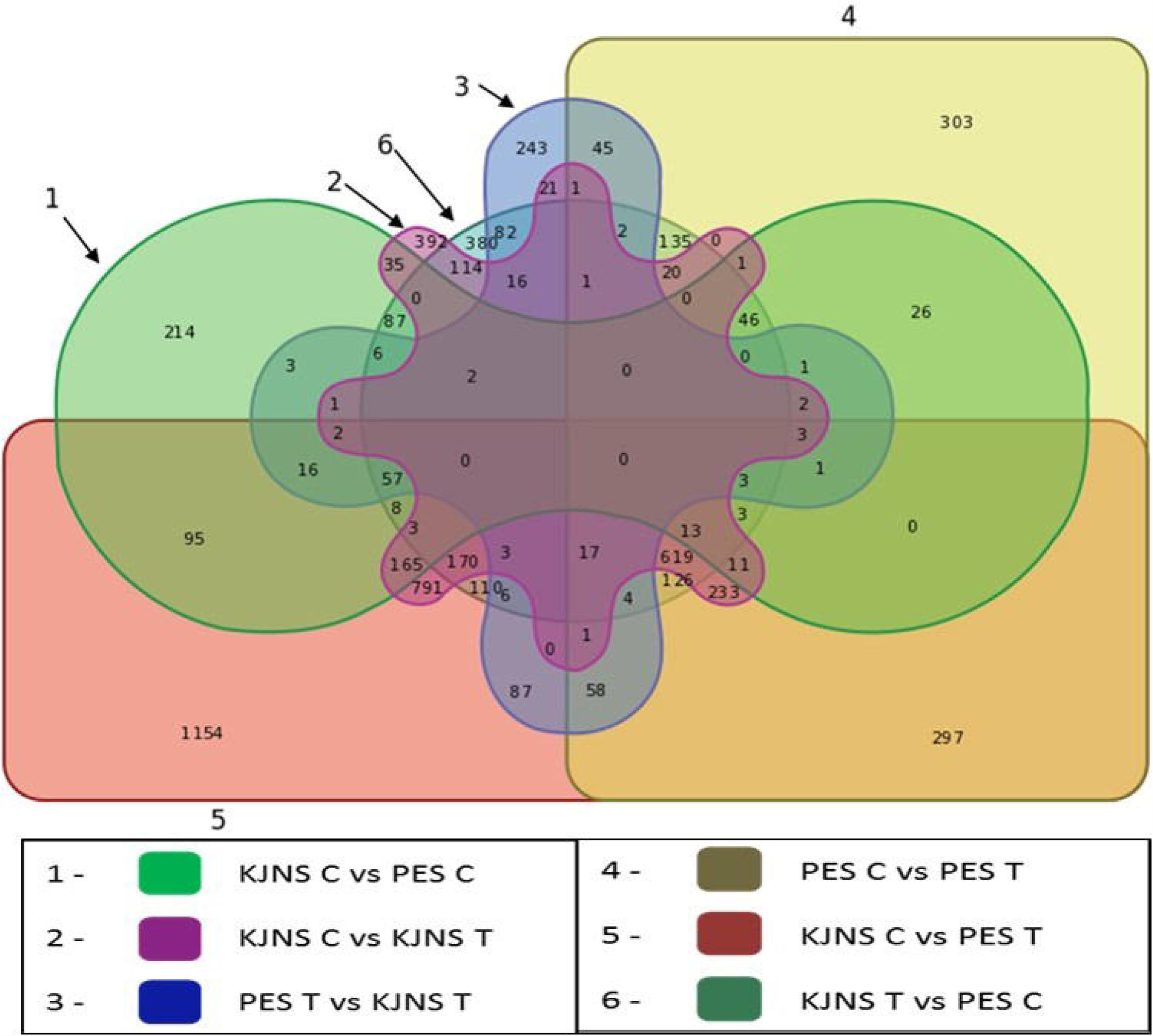
Venn diagram representation of the differentially expressed genes in *E. coracana* transcriptome.

In the present study, we focused on differentially expressed genes in both treatments, i.e., PES-T vs. KJNS-T. Analysis of expression dynamics revealed that only 684 transcripts were differentially expressed under heat stress treatment in both genotypes. Of these, 494 transcripts were upregulated, whereas 190 transcripts were downregulated in PES-T compared to KJNS-T. This result indicated that more transcripts were overexpressed under heat stress in the tolerant genotype (PES-110).

### 3.4 Annotation and functional characterization of DEGs under treatment conditions in both the genotypes

Functional annotation against the NCBI nr protein database of these 684 DEGs was analyzed as mentioned in the material and methods section. In total, 380 (55.56%) DEGs were successfully annotated, whereas 44.44% did not show homology with any protein in the database.

#### 3.4.1 GO classification

To study the putative function of differentially expressed unigenes in PES-T and KJNS-T, sequence homology against GO classification was performed. It revealed that out of 380 annotated DEGs, 196 unigenes were summarized under three main GO categories, including 16 functional groups at level 2 (Fig. 2A). In total, 588 GO assignments were obtained, of which biological processes were the most significant category (38.87%), followed by cellular component (36.73%) and molecular functions (24.40%). In the “biological process” category, 32.36 % of unigenes were involved in the “metabolic process,” 22.01% in the “cellular process,” and 19.09% in the “single-organism process.” The major three groups for the “cellular component” were “cell, membrane, and cell part”, which included 20.55%, 19.52%, and 19.18% of unigenes, respectively. Under “molecular function,” there were only two categories, which included “binding” (51.55%) and “catalytic activity” (48.45%) (Supplementary Table S3, Fig. 2A).

**Fig 2.**
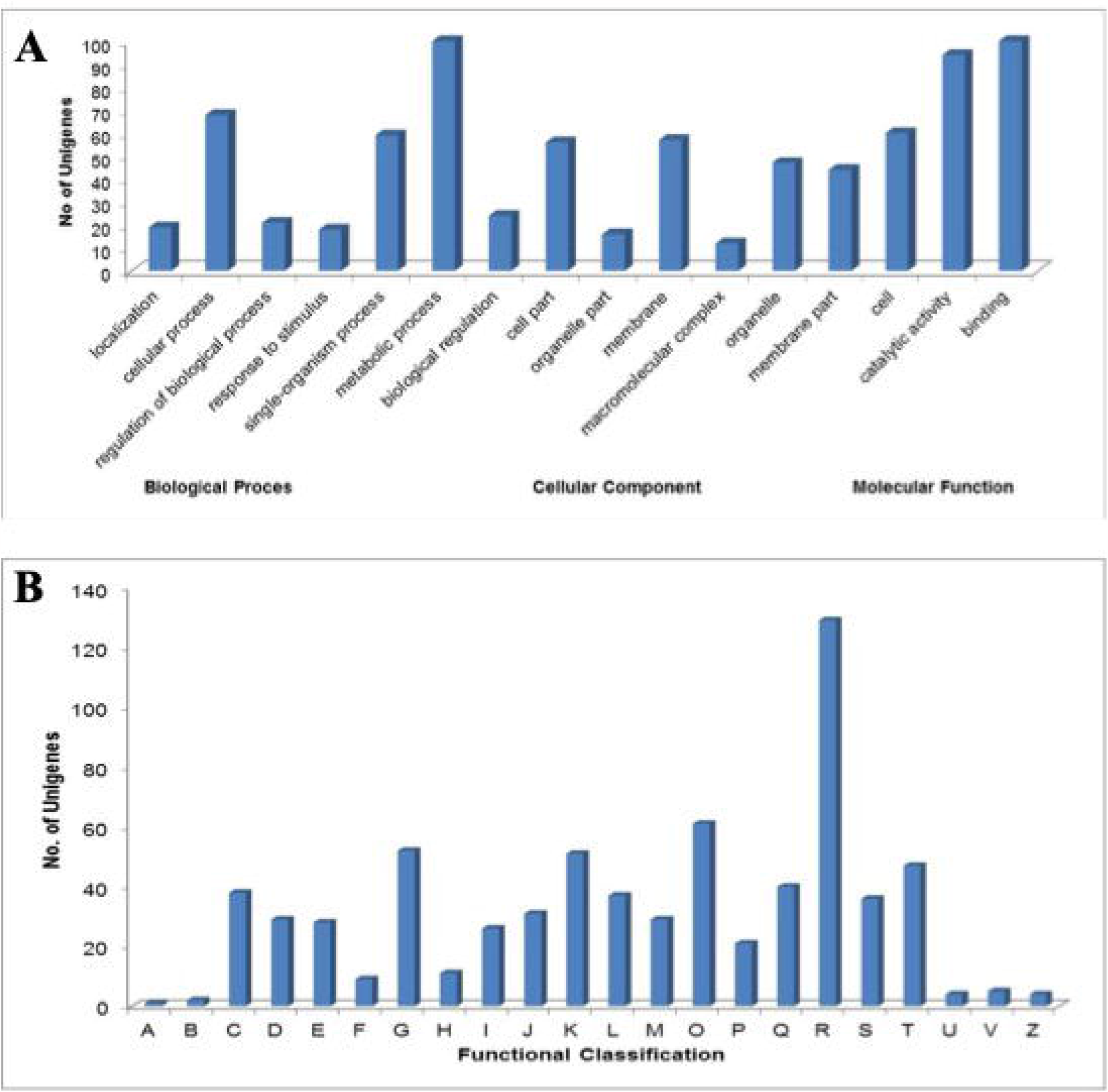
(A) GO classification of *E. coracana* transcriptome and differentially expressed genes between control and salt stress. The Y-axis represents a number of unigenes, and X-axis shows the GO categories. (B) **COG function classification of *E. coracana* transcriptome**. The number of unigenes is reported on Y-axis.

#### 3.4.2 Putative function of unigenes

COG classification of 380 expressed genes indicated that 249 unigenes were clustered into 22 functional categories (Fig. 2B). Some unigenes had multiple COG functions resulting in a total of 691 functional annotations. Out of 22 functional categories, “general functional prediction only” (18.67%) represented the central group, followed by “post-translational modification, protein turnover, chaperone function” (8.82%), “carbohydrate metabolism and transport” (7.53%), “transcription” (7.51%) and “signal transduction” (6.80%).

#### 3.4.3 Pathway analysis

For an improved understanding of the interactions of the putative proteins and biological functions obtained from 684 unigenes, pathway analysis was performed. KEGG orthologous numbers were assigned to 153 unigenes involved in 138 pathways (Supplementary Table S4). In the present study, “Metabolic pathways”, “Biosynthesis of secondary metabolites”, “ribosome”, and “Protein processing in endoplasmic reticulum” were the four largest pathway groups. The 153 KEGG annotated unigenes were categorized into six different categories, including “metabolism (41.02%)”, “genetic information processing (12.88%)”, “environmental information processing (8.81%)”, “cellular processes (5.76%)”, “organismal systems (39%)” and “human diseases (18.31%)”. The largest group comprising 41.02% of unigenes was “metabolism”, with most of the unigenes involved in “general and overview maps” followed by “amino acid metabolism”, “Xenobiotics biodegradation,” and “carbohydrate metabolism”. This suggests that under heat stress, seedlings of finger millet underwent several changes at the molecular level.

#### 3.4.4 Identification of transcription factor (TF)

TF is a vital upstream regulatory protein responding to environmental stresses and developmental processes. The present study classified 39 unigenes, representing 5.70% of the differentially expressed genes between PES-T and KJNS-T, into 16 TF families (Supplementary Table S5). Among these, “HSF type DNA binding” represented the most abundant category comprising 25.64% unigenes. HSF-type DNA binding transcription factor family is a heat shock factor that mediates heat shock gene expression regulation in eukaryotes. Other TF families in our study were “AP2-EREBP” and “NAM” transcription factors representing 10.26% of unigenes.

### 3.5 Common molecular response of finger millet under heat stress

#### 3.5.1 HSPs and sHSPs

It is well known that heat stress generally induces the expression of HSPs in response to different stresses, which is also vital for acclimatization. In the present study, we found that heat stress increased the expression of various HSPs and small heat shock proteins (sHSPs). These were HSP 70, HSP 90, HSP 1, calmodulin binding heat shock protein, HSP 101, HSP 811, HSP 83, and HSP 20. In heat stress samples of both genotypes, all the HSPs have significantly upregulated in our study.

#### 3.5.2 Transcription Factors expressed under heat stress

In the present study, several TFs, including ERF, DREB2A, DREB2B, MYB, AP2, bZIP, WRKY, NAC, DEAD-box RNA helicases, etc., were found to be upregulated. Interestingly, DREB2A TF was found to be up-regulated only in the tolerant genotype (PES-110). We also found several DEGs, like *WRKY* 26, 48, and 7, which were down-regulated in PES-110 treated sample compared to the KJNS-46 sample. HSF30 belonging to the HSFA2 TF family, HSF3 HSFA2e, HSFA2c, and HSBP1 were highly up-regulated in the leaves of thermo-tolerant accession of finger millet (PES-110).

#### 3.5.3 Calcium and Kinases signaling genes and antioxidant enzymes modulated by heat stress

The genes encoding the components of calcium transporting ATPase, a calcium-dependent protein kinase CPK1 adapter protein, calnexin precursor, annexin, calcium/calmodulin mediated signal pathway, calcineurin B, mitogen-activated protein kinases, cell wall-associated protein kinase were induced under heat treatment, in the present study. This suggests that Ca^2+^-mediated signals are essential in response to heat stress treatment in finger millet. Some of the common antioxidants, like, APX, DHAR, SOD, catalase, glutathione-s-transferase, and peroxidase, were up-regulated, suggesting these genes may have a fundamental role in response to heat stress. Genes related to cell rescue, including *stress-induced proteins* and *galactinol synthase*, also exhibited a change in expression. Some genes such as metabolism-associated enzymes, 3-hydroxy-3-methylglutaryl coenzyme A (acetyl-CoA pathway), development-related enzymes, and ripening-regulated protein also changed expression patterns in response to heat stress in the present study. Further, validation and deep investigation in this direction are needed to understand better the molecular mechanism involved.

#### 3.5.4 Metabolism

Galactinol synthase transcript level was significantly increased in response to heat stress in finger millet. In addition, carbohydrate metabolism genes include *invertase, UDP-glucose dehydrogenase, 6-phosphate dehydrogenase, sucrose synthase*, and *trehalose-6-phosphate synthase/phosphatase*. These enzymes are needed for sugar metabolism and were downregulated by heat stress treatment in the present study. In addition, heat stress inhibited lipid metabolism, affecting L-asparaginase and lipoxygenase.

### 3.6 Validation using qRT-PCR

To validate the result of RNA sequencing analysis, we carried out the qRT-PCR analysis of twelve genes, as mentioned in the material and methods. The result showed that all the verified genes followed the same expression pattern as observed in the Illumina sequencing, validating the transcriptome results (Fig. 3). Moreover, the fold-changes obtained by qRT-PCR were more significant than those obtained by DEGs, which have also been reported in other studies and attributed to the essentially different of algorithms and sensitivity between the two techniques (Goyal et al. 2016a, b). This result indicated that the method used in the study to determine DEGs were valid.

**Fig 3.**
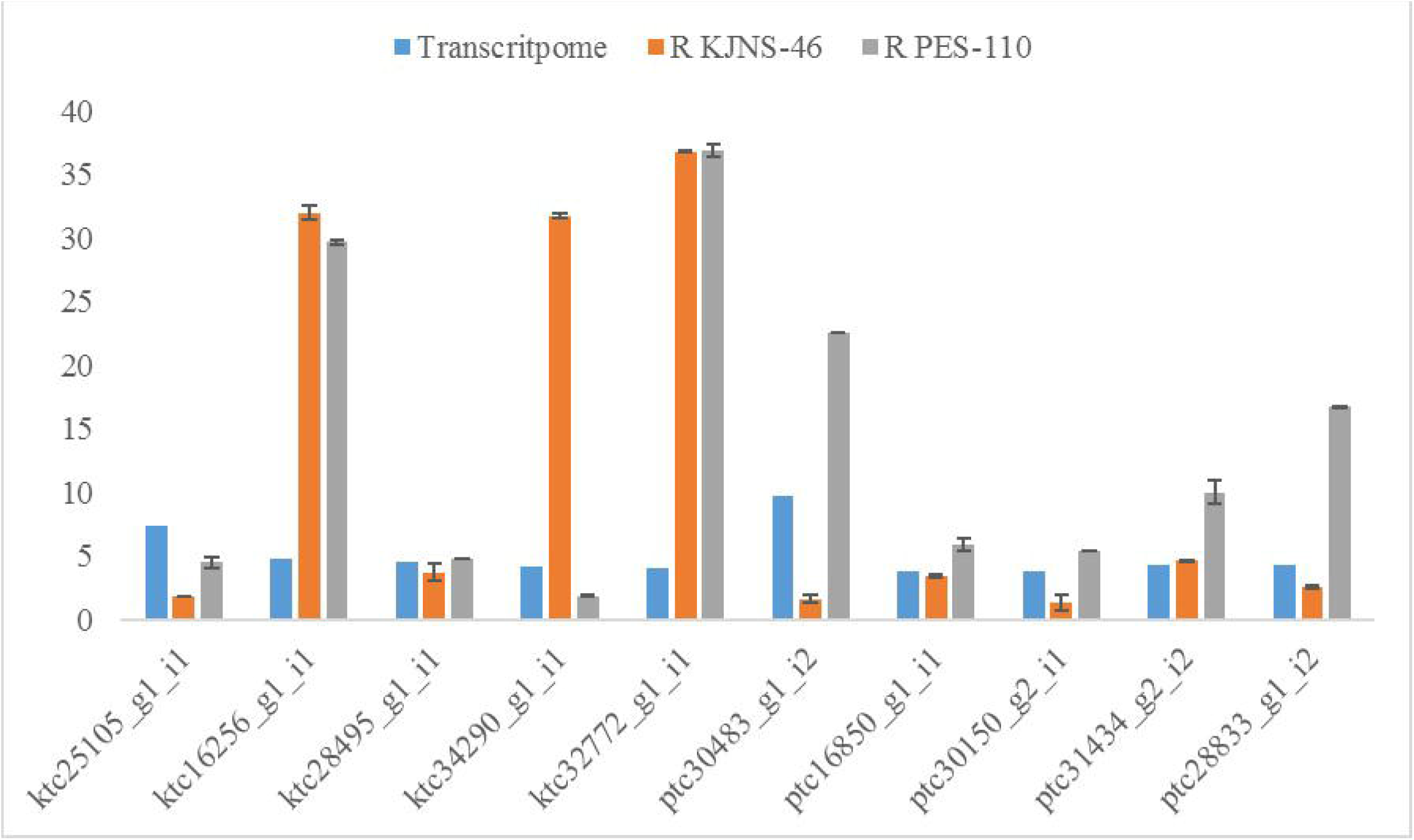
Real-time validation of some selected genes of *E. coracana* transcriptome.

## 4. Discussion

Several independent and dependent biological pathways are regulated to confer tolerance towards to heat stress. Due to climate change, exploring the complex molecular mechanism of plant response under heat stress has become a central subject of agricultural significance (Baniwal et al. 2014). Transcriptome sequencing has been recognized as a rudimentary technique that provides a genome-wide, quantitative view of how changes in environmental conditions can affect the transcription pattern of genes. In their response to various abiotic stresses, studying contrasting genotypes provides a great chance to understand the molecular basis of variability. It will help us decipher the interactions of the complex responses at the transcriptome level. Therefore, this study was designed to question the difference at the molecular level in contrasting finger millet genotypes (PES-110 and KJNS-46) in response to high-temperature stress. This will provide a clear picture of the genes regulating heat stress tolerance or susceptibility in the contrasting genotypes.

The differential expression analysis of RNA-seq data in ‘KJNS-46’ and ‘PES-110’ showed that many genes expressed in response to heat treatment of ‘PES-110’ were higher than in ‘KJNS-46’ (Fig 1). This result indicated that under stress, ‘PES-110’ could better increase transcriptional regulation than ‘KJNS-46’. Many previous studies also discovered an evident expressional divergence between heat-tolerant and sensitive strains of pepper, jujube, and *P. haitanensis* in response to high-temperature stress (Li et al. 2015, Singh et al. 1973, Wang et al. 2018). Heat stress-responsive genes and proteins in plants can be divided into two groups. Signaling components are the first group, including protein kinases and transcription factors. On the other hand, functional genes such as heat shock proteins (HSPs) and catalase (CAT) belong to the second group. The major HSPs are divided into five structurally distinct classes: Hsp100, Hsp90, Hsp70, Hsp60, and small HSPs (sHSPs) (Bharti et al. 2002). The induction of HSPs expression is one of the common heat-responsive mechanisms in all organisms (Kotak et al. 2007; Queitsch et al. 2000; Yamada et al. 2007). Myouga et al. (2006) reported that the HSP101 homolog conferred thermo-tolerance to chloroplasts during heat stress. It is well known that HSP101 interacted with the sHSPs chaperone system to resolubilize protein aggregates after heat stress. Thereby improving tolerance to heat stress in plants such as *Arabidopsis* and maturing tomato pollen grains (Lee et al. 2005; Pressman et al. 2007; Frank et al. 2009). Heat stress also induces small proteins called sHSPs (Small heat shock proteins), having monomeric masses between 12 to 43 kDa. Combined with other HSPs, they prevent the aggregation of proteins and play an essential role in their refolding. A strong correlation between sHSPs accumulation and plant tolerance to heat stress has been proved before (Pressman et al. 2007). In this study, most HSP genes, including HSP90, HSP70, and sHSPs, were upregulated in response to heat stress, which is in correlation with other studies as mentioned above (Myouga et al. 2006; Lee et al. 2005; Pressman et al. 2007; Wang et al. 2004). Thus, the upregulated HSP genes play essential roles in heat tolerance in ragi.

Abiotic stresses activate many transcription factors that have been reported to act as fundamental factors in regulating the expression of genes (Wahid et al. 2007; Golldack et al. 2011). Therefore, identifying and characterizing TFs involved in heat stress response is key in revealing the molecular mechanisms. This study identified the activation of differentially expressed transcription factors, including ERF, WRKY, MYB, NAC, DREB, bZIP, and HSF, in response to heat stress (Table S5). Basic Leucine Zipper (bZIP) transcription factors have been reported to be involved in broad functions in response to various biotic and abiotic stresses (Jacoby et al. 2002). NAC (NAM, ATAF1/2, CUC2) transcription factors family is known to activate the expression of “early responsive to dehydration stress 1 (ERD1),” which is predominantly induced by abiotic stress in guard cells (Tran et al. 2004). Guo et al. (2015) reported that overexpressing *TaNAC2L* from wheat improved acquired heat tolerance and activated the expression of heat-related genes in transgenic *Arabidopsis* plants. MYB gene function in ABA signaling and jasmonic acid-related gene expression, indicating that they are involved in crosstalk between abiotic and biotic stress responses. Expression of DREB 2A and DREB 2B in *Arabidopsis* increased transiently within one hour of heat stress. In contrast, DREB 2 expression was up-regulated even 12 hours after the heat treatment, as reported earlier (Sakuma et al. 2006). bZIP, MYB, DREB, and NAC TFs were upregulated in the present study, which is in correlation to the above-mentioned study, proving how TFs are essential in conferring tolerance towards heat stress. Several pieces of evidence indicate that several WRKY genes also regulate the response to various abiotic stresses, including heat (Miller et al. 2008). In the present study, some *WRKY* genes were downregulated, per the previous reports in grapes and rice (Schain et al. 2016; Liu et al. 2012). The exact role of all the remaining *WRKY* genes downregulated in the present study is not fully understood and needs to be elucidated. HSFs play indispensable roles in basal and acquired thermo-tolerance through binding to *cis*-acting regulatory elements called heat shock elements (HSEs) in the promoter region of the *HSPs* gene. Earlier reports on mature tomato microspores, grape, rice, and *Arabidopsis* showed enhanced expression of either TFs in response to heat stress, which is in regulation with our study (Liu et al. 2012; Schain et al. 2016; Pressman et al. 2007; Prandl et al. 1998).

Abiotic stress, especially heat stress, also activates several regulatory networks, which include signal transduction pathways to generate a series of innate defensive reactions (Yan et al., 2016). Calcium signal is vital in response to heat stress, which causes a rise in Ca^2+^ levels in many plants. In wheat, several genes related to calcium signal pathways were heat regulated, including annexin, calcium-binding proteins (CBPs), calcium-dependent protein kinases (CDPKs), voltage-gated calcium channel activity, Ca^2+^ binding protein EF-hand, CBL, CIPK (Qin et al. 2008). In the present research, some genes were highly homologous to calcium signaling genes, including CaM, CDPK, CBK, and CBL, as reported in grapes, spinach, and pepper (Liu et al. 2012; Li et al. 2015; Yan et al. 2016).

## 5. Conclusion

The data here provides genome-wide expression profiles of contrasting finger millet genotypes under heat stress in contrasting genomes. Illumina Hiseq 2000 and qRT-PCR analysis were used to identify heat stress-responsive genes representing classic and heat stress-responsive and thermo-tolerance mechanisms. The present study highlighted the noteworthy contribution and essential roles of transcriptional control in stress responses in finger millet. Control and heat stress responses displayed differential gene expression changes. We found that 684 DEGs were responsive under heat stress treatment in both genotypes (KJNS-T vs. PES-T). The number of heat stress up-regulated genes was almost thrice that of down-regulated genes in PES-T compared to KJNS-T. The heat-responsive genes identified in this study belong to many important factors and biological pathways, including those for cell rescue (i.e., antioxidant enzymes), protein fate (i.e., HSPs), primary and secondary metabolism, transcription factors, signal transduction, hormone, calcium, and kinase signaling and development (Fig. 4). Of particular interest, HSPs especially sHSPs, APX, and galactinol synthase, may be essential to thermo-tolerance. Identified genes will provide rational candidates for germplasm screening to enhance heat tolerance in finger millet.

**Fig 4.**
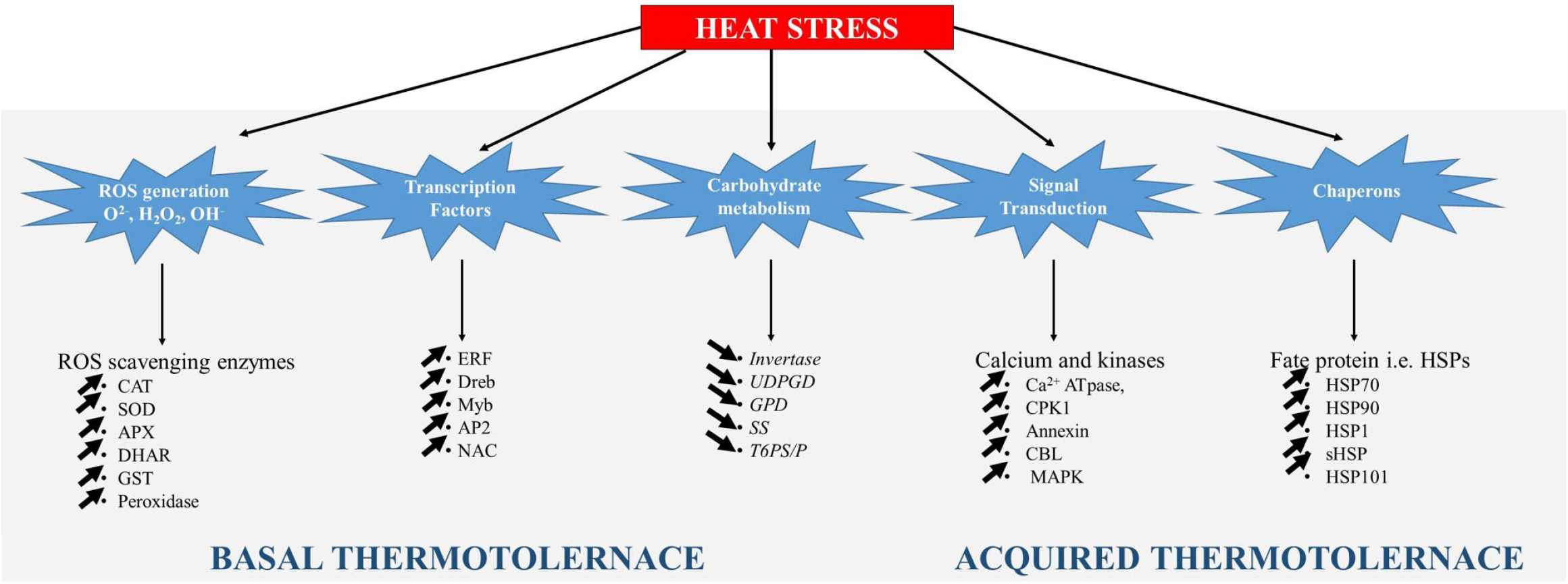
Schematic representation of the proposed model revealing the possible mechanism in response to heat stress tolerance in finger millet.

These results provide novel insight into the finger millet leaf response to heat stress and have significant implications for further gene function annotation and molecular breeding studies. The information from this study can be used to develop new plant types that can adapt and thrive well under high temperatures predicted for the future. The putative functional unigenes identified in the present study can provide leads for future investigations. An improved understanding of their precise roles, functions, and contribution to heat stress tolerance will enable us to employ them for improving tolerance to heat stress. Identifying the specific gene set in the heat stress regime and elucidating the complete mechanisms of heat stress response will contribute to fine-scale control in future breeding programs and predict the response of ragi to changes in climatic conditions.

## Acknowledgment

This financial support from the Department of Biotechnology, New Delhi, under the project “Phenomics and Genomics of Ragi (*Eleusine coracana*),” is acknowledged. The authors are grateful to Prof. Y.A. Nanja Reddy, Project Coordinator, AICP on Small Millets, UAS, GKVK Campus, Bangalore, for providing seeds of KJNS-46 and PES-110 and valuable suggestions.

## Supplementary Tables

**Supplementary Table 1 Primers used in the study**.

**Supplementary Table 2 Sequence annotation of *E. coracana* transcriptome**. (A) KJNS-46; (B) PES-110.

**Supplementary Table 3 GO classification of 684 differentially expressed unigenes**.

**Supplementary Table 4 KEGG classification of 684 differentially expressed unigenes**.

**Supplementary Table 5 Distribution of Transcription factor family of 684 differentially expressed unigenes**.

## Supplementary Figure

**Supplementary Fig 1 Transcriptome assembly information**. (a) Length distribution of fully assembled transcripts. (b) Proteome analysis of total transcripts of *E. coracana*.

