## Supplementary figures and images for "Comparative transcriptome analysis in contrasting finger millet (*Eleusine coracana* (L.) Gaertn) genotypes for heat stress"

### Fig S1.jpg

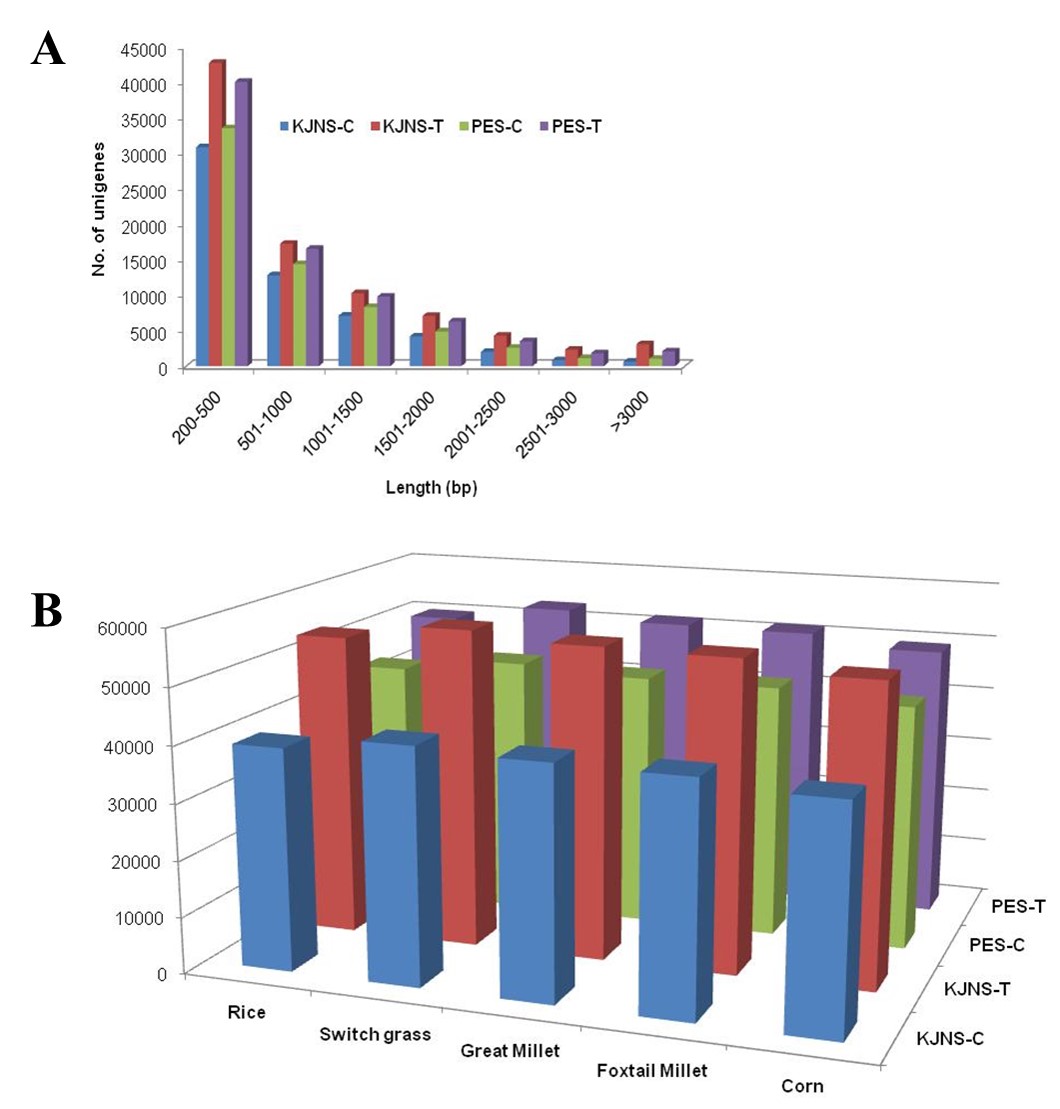
